# Aging-related cerebral microvascular changes visualized using Ultrasound Localization Microscopy in the living mouse

**DOI:** 10.1101/2021.06.04.447141

**Authors:** Matthew R. Lowerison, Nathiya Chandra Sekaran, Wei Zhang, Zhijie Dong, Xi Chen, Daniel A. Llano, Pengfei Song

## Abstract

Aging-related cognitive decline is an emerging health crisis; however, no established unifying mechanism has been identified for the cognitive impairments seen in an aging population. A vascular hypothesis of cognitive decline has been proposed but is difficult to test given the contradictory radiologic needs of high-fidelity microvascular imaging resolution and a broad and deep brain imaging field of view. Super-resolution ultrasound localization microscopy (ULM) offers a potential solution by exploiting circulating microbubbles to achieve a vascular resolution approaching the capillary scale without sacrificing imaging depth. In this report, we apply ULM imaging to a mouse model of aging and quantify differences in cerebral vascularity, blood velocity, and vessel tortuosity across several brain regions. We found significant decreases in blood velocity, and significant increases in vascular tortuosity, across all brain regions in the aged cohort, and significant decreases in blood volume in the cortex. These data provide the first-ever measurements of subcortical microvascular dynamics *in vivo* and reveal that aging has a major impact on these measurements.

## Introduction

The U.S. population is aging, with the number of adults over the age of 65 expected to nearly double by the year 2050 (1). This aging population is more vulnerable to cognitive impairment and dementia (2–4), which represents an emerging public health crisis with wide-reaching implications for quality of life and the economic burden of care (5). However, aging-related cognitive decline remains a controversial area of research, with no established unifying mechanism identified for the cognitive and memory impairments seen in an aging population. Two of the most commonly observed pathological findings in the aged brain, both in human and in animal models, is decreased microvascular density and increased vessel tortuosity (6–12). These findings imply that there is a relationship between compromised cerebral blood flow and cognitive impairment, where there are paralleled deteriorations in both small vessel function and cognitive ability. This relationship is reinforced by overlapping epidemiological risk factors (13); both late-stage cognitive decline and vascular disease are associated with diabetes (14), obesity (15), and hypertension (16). Furthermore, clinical evidence suggests that microvascular changes accelerate clinical decline in cognitive impairment (17,18), and decreased macroscopic cerebral blood flow is associated with worsened cognitive performance in both normal and pathological states (19–21).

However, the vascular hypothesis of cognitive decline remains difficult to test within a clinical context, given the contradictory radiologic needs of high-fidelity microvascular imaging resolution and a broad and deep brain imaging field of view. The aging-associated decreases in cerebral microvascular density are heterogenous across brain regions (12,22,23), implying a selective vascular vulnerability that requires an imaging modality that provides a broad field of view, high depth of penetration, and high spatial resolution. Traditional approaches in animal studies include histological analysis of the microvasculature, which does not provide information about the dynamics of blood flow. Multiphoton imaging may be applied to the surface of the brain to measure dynamics but cannot be applied to deep brain regions without the use of tissue-destructing prisms, periscopes, or fibers. Macroscopic blood flow measurement techniques such as single-photon emission computed tomography (SPECT) or MRI-based methods do not provide adequate spatial resolution to measure microscopic blood flow dynamics.

A recently proposed ultrasound-based super-resolution imaging modality, ultrasound localization microscopy (ULM) (24–27), can potentially address the need for microvascular imaging fidelity simultaneously with deep brain imaging. ULM processing leverages an intravascular injection of clinically used microbubble contrast agent to function as an imaging point target to break the diffraction limit of ultrasound. This technique can improve the vascular imaging resolution by a factor of about ten (28) over conventional ultrasound imaging, but does not sacrifice the imaging penetration depth, thereby side-stepping the classical compromise between imaging resolution and imaging depth. This improvement results in the ability to produce microvascular network reconstructions, potentially down to the capillary scale, that can cover an entire cross-section of the brain. The technique also retains the non-invasiveness and safety profile of conventional contrast-enhanced ultrasound imaging.

In this report, we apply super-resolution ULM imaging to a mouse model of aging and quantify differences in cerebral vascularity, blood velocity, and vessel tortuosity across several brain regions. Given the aging-related changes that occur in the peripheral and central auditory system (29,30), we focused on central auditory structures and measured peripheral auditory function. A total of 16 young (average age in weeks = 27.7) and 11 aged (average age in weeks = 106.8) mice of either sex were used in this study. Ultrafast (1000 Hz frame rate) contrast-enhanced ultrasound data were acquired using a Verasonics Vantage 256 ultrasound system and a L36-16vX high-frequency linear array transducer through a cranial window. ULM images were reconstructed at a 4.928 µm isotropic axial/lateral resolution. We found a reduction in cortical vascularity in the aged mouse group, and significant decreases in blood velocity and increases in tortuosity across multiple anatomical regions in comparison to the young mouse group. These data provide the first-ever measurements of subcortical microvascular dynamics *in vivo* and reveal that aging has a major impact on these measurements.

## Materials and Methods

### Animal model

All procedures on mice presented in this manuscript were approved by the Institutional Animal Care and Use Committee (IACUC) at the University of Illinois Urbana-Champaign. Mice were housed in an animal care facility approved by the Association for Assessment and Accreditation of Laboratory Animal Care. CBA/CaJ mice were bred in-house. A total of 16 young (average age in weeks = 27.7, max = 46.8, min = 11.6) and 11 aged (average age in weeks = 106.8, max = 126.9, min = 90.6) CBA/CaJ mice of either sex were used in this study.

### Hearing threshold testing

For auditory brainstem responses, animals were anesthetized with intraperitoneal ketamine hydrochloride (100 mg/kg) and xylazine (3 mg/kg) before the insertion of three subdermal electrodes: one at the vertex, one behind the right ear, and a reference electrode placed behind the left ear. Stimuli were presented using a System 3, ES1 free-field speaker (Tucker-Davis Technologies), with waveforms generated by SigGen software. The output of the speaker was calibrated at the relevant frequencies, using a microphone (Model 377A06; PCB) and a preamplifier (SV 12L; Svantek). The stimulus was presented for 5 ms (3 ms flat with 1 ms for both rise and fall times), at a rate of 21 Hz, with a 45 ms analysis window. Raw potentials were obtained with a RA4LI headstage, RA16PA preamp, and RA16 Medusa Base station (Tucker-Davis Technologies), and filtered between 100 and 5000 Hz. Significant deflections, assessed via visual inspection, within 10 ms after the onset of the stimulus were deemed a response.

### Animal preparation for ultrasound imaging

Mouse anesthesia was induced using a gas induction chamber supplied with 4% isoflurane mixed with medical oxygen, and then mice were placed in a stereotaxic frame with nose cone supplying 2% isoflurane with oxygen for maintenance. Lidocaine (1%) was intradermally injected in the scalp to supplement anesthesia. Ear bars were used to secure the mouse head to the stereotaxic imaging stage. The scalp of the mouse was removed, and a cranial window was opened on the left side of the skull using a rotary Dremel tool, starting at the sagittal suture and moving laterally to expose the auditory cortex. The tail vein of the mouse was cannulated with a 30-gauge catheter and vessel patency was confirmed with a 0.1 mL injection of sterile saline.

### Ultrasound imaging

A Verasonics Vantage 256 system (Verasonics Inc., Kirkland, WA) was used for all ultrasound imaging in this study. A L35-16vX transducer (Verasonics) was secured via a 3D-printed transducer holder to a translation motor (VT-80 linear stage, Physik Instrumente, Auburn, MA) that was connected to the stereotaxic imaging frame. The transducer was then positioned above bregma, via visual inspection, oriented to produce a coronal anatomical section of the brain. Acoustic contact gel was applied directly to the surface of the mouse brain and adjacent skull, and the transducer was lowered into position. Once acoustic coupling was confirmed, the motorized stage was adjusted at 0.1mm increments caudally to find an appropriate imaging plane which contained the auditory cortex and auditory thalamus. This was generally 3.0-3.5mm caudal to bregma. The imaging field of view was then adjusted to cover the entire half of the mouse brain in this anatomical position, and a 400-frame scouting dataset was acquired and processed to produce a power Doppler image as in (31) to confirm transducer placement over the anatomy of interest (**Figure 2C**). A 50uL bolus of a clinically available ultrasound contrast agent (DEFINITY®, Lantheus Medical Imaging, Inc.) was injected into the mouse just prior to starting ULM image acquisition. Imaging was performed with a center frequency of 20 MHz, using 9-angle plane wave compounding (1-degree increments) with a post-compounding frame rate of 1,000 Hz. Ultrasound data was saved as in-phase quadrature (IQ) datasets for off-line processing in MATLAB (The MathWorks, Natick, MA; version R2019a). A fresh 50uL bolus of microbubbles was injected after every 10 imaging acquisitions (1,600 frames per acquisition). A total of 80 acquisitions (128,000 frames, or 128 seconds of data) was acquired for this imaging plane. The transducer was then moved 0.5mm in the rostral direction, and the contrast microbubble imaging procedure was repeated for another 80 acquisitions at this second imaging plane.

### Ultrasound signal processing and super-resolution reconstruction

The isolated microbubble signals were extracted from the contrast-enhanced IQ data by applying a spatiotemporal singular value decomposition (SVD)-based clutter filter (32–35). Briefly, each IQ dataset was first reshaped into a columnized 2D Casorati matrix and an SVD decomposition was performed to reveal the singular values of the data. A low-order singular value threshold was determined adaptively (36) to filter out tissue signal, which typically zeroed out the first 10-20 singular values. An inverse SVD was then performed, and the data was reshaped into the original data size. A noise-equalization profile (37) was then applied to equalize the microbubble signal intensity through the entire depth of imaging.

An isolated microbubble signal was manually identified, and a multivariate Gaussian function was empirically fit to the axial and lateral dimensions to represent the point-spread function (PSF) of the system. A microbubble separation filter (34) was then applied to the SVD-filtered IQ dataset to separate it into data subsets that have more spatially sparse microbubble distributions. Each IQ data subset was then spatially interpolated to an isotropic 4.928 µm axial/lateral resolution using 2D spline interpolation (38). A normalized 2D cross-correlation was then performed with the empirical PSF to localize microbubbles on every frame. Pixels with a low cross-correlation coefficient were excluded via a threshold (33,34,39), and microbubble centroids were localized with the MATLAB built-in “imregionalmax.m” function. Frame-to-frame microbubble centroid pairing and trajectory estimation was performed using the uTrack algorithm (40). A minimum microbubble trajectory length of 20 frames (i.e., 20 ms) was applied to the super-resolution reconstructions presented in this study. Motion was corrected for each ULM accumulation by using a 2D normalized cross-correlation with the first processed acquisition.

### Ultrasound image analysis

Brain anatomical regions (auditory cortex, auditory thalamus, entorhinal cortex, hippocampus, superior colliculus, and visual cortex) were segmented by manually placing Bezier control vertices on the border of the region of interest (ROI) and by interpolating with Hobby’s algorithm (41). Brain vascularity was calculated by binarizing the super-resolution vessel maps to determine the percentage of cross-section that was perfused. Blood vessel velocity was determined for every microbubble track directly from the frame-to-frame displacement of detected microbubble centroids. The sum of angles metric, a measure of vascular tortuosity, was calculated for every microbubble trajectory using the algorithm described by Shelton *et al*. (42).

### Statistics

All statistical analysis was performed in the R programming language (43), and all graphs were generated using the ggplot2 package (44). A two-way analysis of variance (ANOVA) was applied to test for statistical significance between young and aged mouse brain anatomy with a Tukey’s honestly significant difference test applied as a post-test. Auditory brain responses and mouse weights were analyzed using a Mann Whitney U test. A corrected p < 0.05 was considered as statistically significant.

## Results

### Hearing threshold is substantially impaired in the aged mouse group

The hearing threshold for each mouse was determined by stimulus-evoked auditory brainstem responses (ABR) and is presented in **Figure 1**. The young mouse group demonstrated a wide range of high-frequency responses and yielded a hearing threshold level (average of 37 dB SPL) which similar to that previously reported for CBA/CaJ mice of that age range (29). The aged mouse group had a severely impaired hearing threshold (average 76.8 dB SPL), as previously-reported (45,46). A Mann Whitney U test confirmed that these differences were statistically significant (p < 0.001). The body weight of the two cohorts showed a trend toward heavier aged mice (**Figure 1B**), but this difference was not significant.

**Figure 1.**
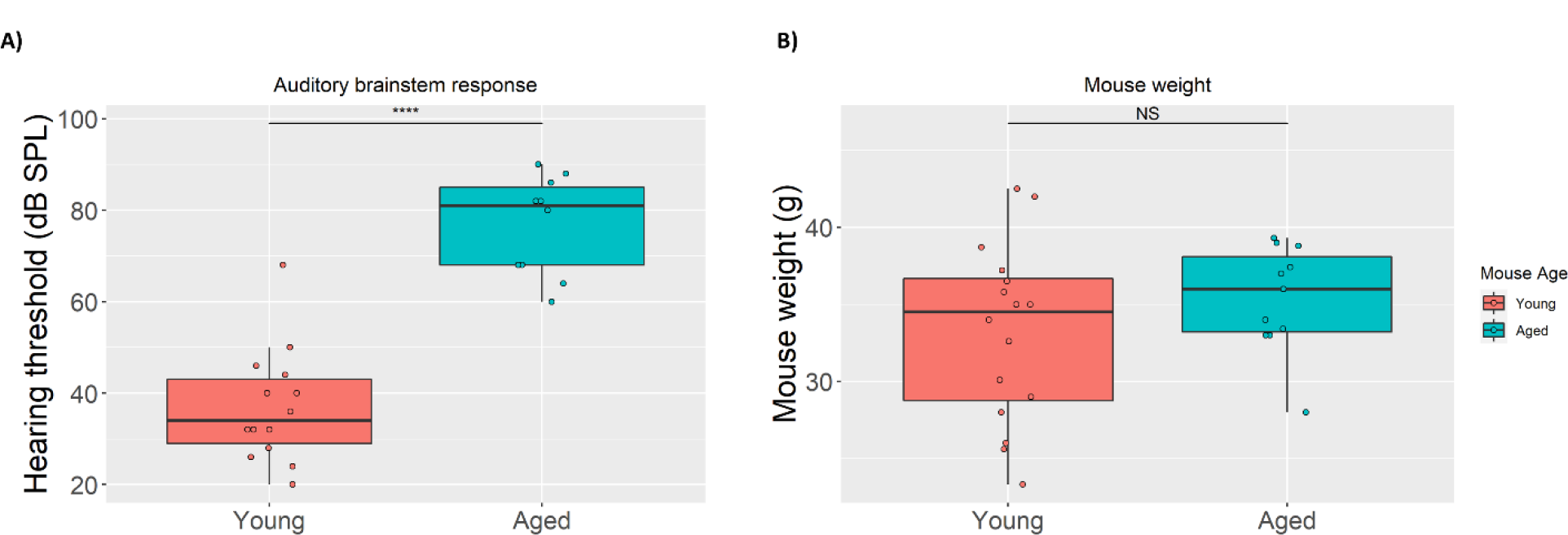
Auditory brainstem response for young and aged mice. (**A**) Auditory thresholds were determined by stimulus-evoked auditory brainstem responses. N = 14 for the young mouse group (average age 6 months) and N = 10 for the aged group (average age 20 months). Statistical significance was determined using a Mann Whitney U test (p < 0.001). Data are presented as boxplots (line at median, box surrounding interquartile range, whiskers ranging from maximum to minimum excluding outliers). Raw values are plotted as individual datapoints. (**B**) The body weight of the two cohorts were not significantly different.

### Ultrasound imaging through cranial window allows simultaneous observation of auditory cortex and auditory thalamus

The unilateral cranial window opened in this mouse cohort extended from the sagittal suture to the side of the skull and was centered around 3 mm caudal from bregma (**Figure 2A**). Extending the cranial window beyond the sagittal midline of the skull was avoided to prevent damage to the sagittal sinus, as this large blood vessel is sensitive to puncture from the Dremel tool used to remove the skull. This placement exposed the hemisphere of the mouse brain that contained both the auditory cortex and the auditory thalamus. Elevational tissue motion was minimal after securing the mouse via ear bars to the stereotaxic frame (**Figure 2B**), however some respiratory motion was still present in the acquired contrast-enhanced IQ dataset. Given that this motion was predominantly in-plane, it could be accounted for via 2D normalized cross-correlation. An example B-mode and power Doppler image (**Figure 2C**) demonstrate the relevant anatomies of interest, with imaging penetration extending to the bottom of braincase. The diffraction-limited power Doppler image, processed as in (31), reveals highly vascularized brain tissue with an evident hierarchical vascular branching morphology, which added in the identification of the anatomies of interest. The corresponding Allen Brain Atlas (47) coronel section (**Figure 2D**) with the ROIs highlighted shows a good correspondence to the ultrasound imaging field of view.

**Figure 2.**
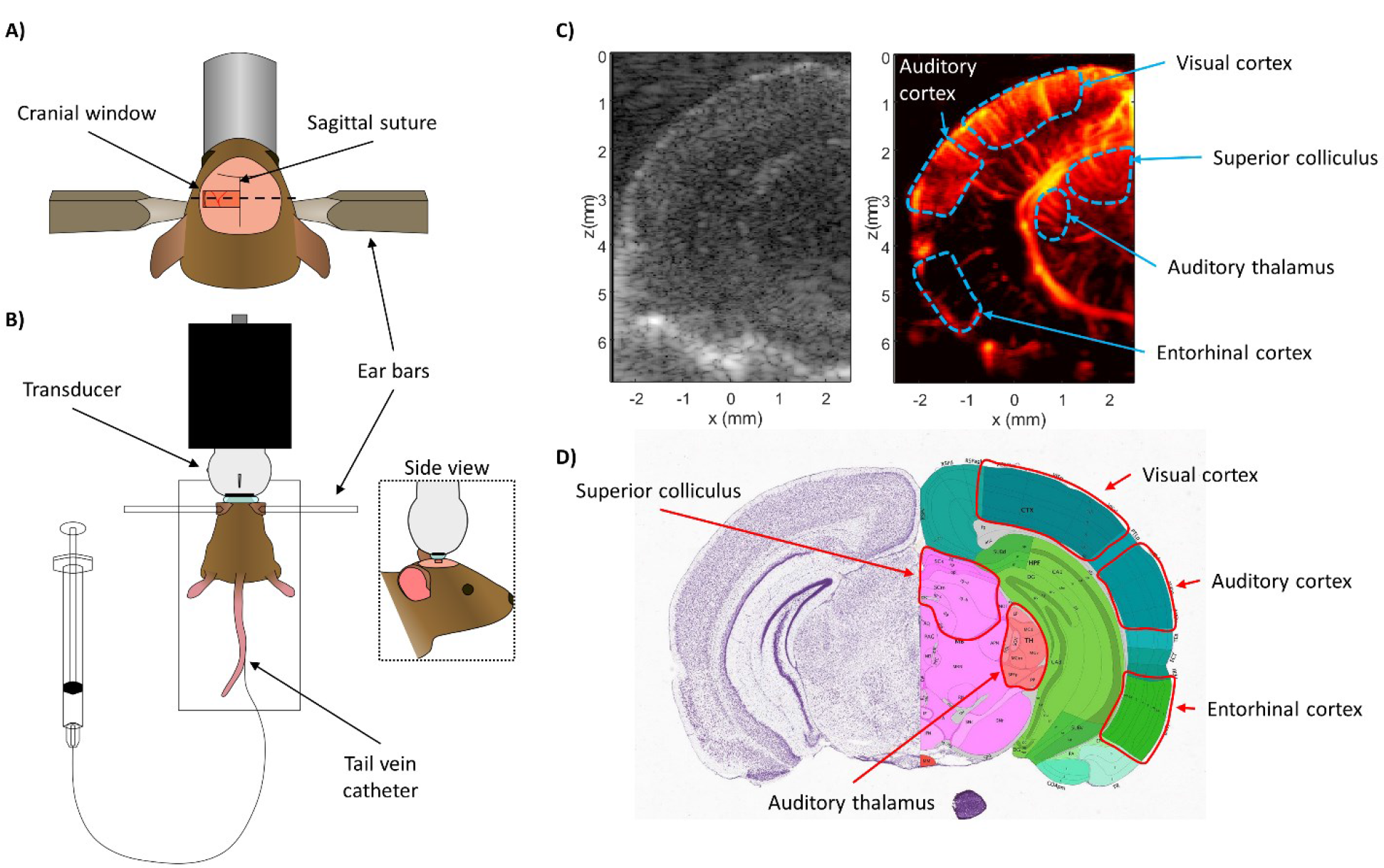
Ultrasound imaging acquisition through cranial window. (**A**) A rectangular cranial window was opened in the mouse skull spanning from the sagittal suture to the lateral side of the skull, centered approximately 3mm away from bregma. This exposed the auditory cortex and auditory thalamus for ultrasound imaging. (**B**) Diagrammatic example of the stereotaxic imaging frame. The mouse was positioned on a heated pad and the skull was secured to the stage using ear bars. The high-frequency ultrasound transducer was connected to a linear translation motor via a custom 3D printed holder. Tail-vein catheterization permitted controlled and repeatable bolus injections of microbubble contrast agent. (**C**) Representative B-mode and power Doppler (no contrast) images from the first imaging plane. The ROIs used in this study (auditory cortex, auditory thalamus, entorhinal cortex, superior colliculus, and visual cortex) are outlined in blue. (**D**) Mouse coronal section taken from the Allen Brain atlas (https://mouse.brain-map.org/static/atlas) shows good correspondence between the features seen in the B-mode and power Doppler image with the anatomies of interest.

### ULM processing permitted high-fidelity vascular mapping through the whole brain depth

Ultrafast planewave imaging of the mouse brain following contrast microbubble injection demonstrated a highly enhancing vascular bed along with the strong acoustic backscatter from tissue (**Figure 3A**). The flowing microbubble signal could be extracted from the tissue background by applying SVD clutter filtering (**Figure 3B**), and the microbubble signal intensity needed to be normalized using a noise equalization profile (37) before any upstream processing. This image processing yielded isolated microbubble signal at a high concentration, which was split into sparser datasets using a microbubble separation filter (34). The microbubbles in these sparser subsets could then be localized by detecting their centroids (**Figure 3C**), producing centroid coordinate vectors that can serve as input into the uTrack algorithm for pairing and tracking. Super-resolution ULM images from each data subset (**Figure 3D**) demonstrated the reconstruction of different vascular features, which depend on the flow direction, velocity, and decorrelation of the microbubble signal. The combination of each of these ULM subset reconstructions resulted in an accumulation image (**Figure 3E**) with a high degree of vascular perfusion given the relatively short imaging period (1,600 frames, or 1.6 seconds of acquisition). The final ULM reconstruction used 80 of these accumulation maps (128 seconds of data) to ensure that the majority of the microvascular bed had been perfused with microbubbles (48–50).

**Figure 3.**
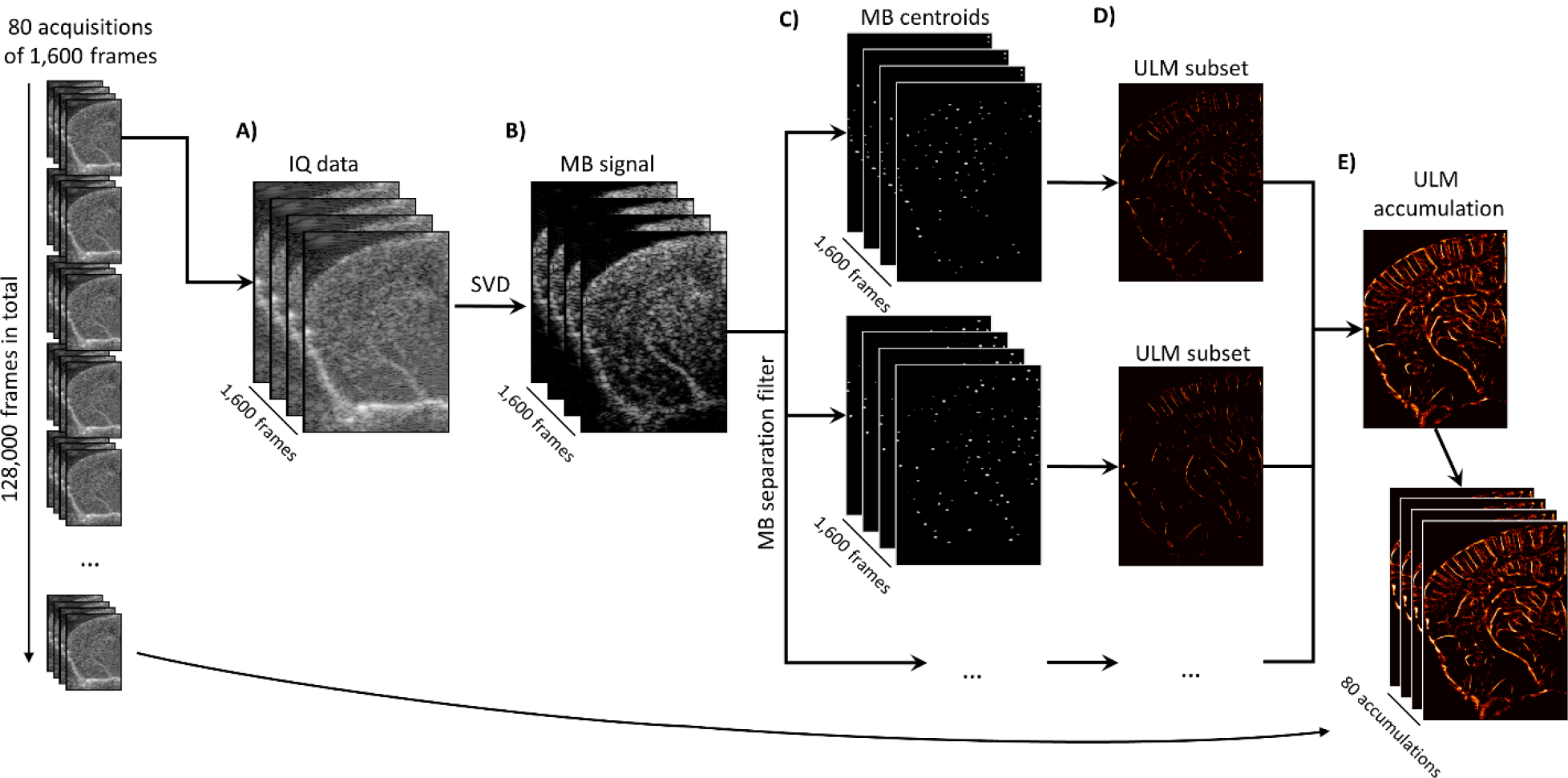
Super-resolution reconstruction workflow. (**A**) Each 1,600 frame IQ data acquisition was represented as a 3D matrix of stacked imaging frames. This was then reshaped into a Casorati matrix and an (**B**) SVD filter was applied to extract the moving microbubble signal from the highly spatiotemporally coherent tissue background. (**C**) A microbubble separation filter was then applied to split the high concentration microbubble dataset into sparser subsets, each of which were independently processed. The microbubble locations were detected on each subset using a 2D normalized cross-correlation with an empirically determined PSF function. (**D**) The uTrack algorithm was then applied on detected centroids to pair microbubbles and estimate trajectories. (**E**) The final reconstruction for this 1,600 frame dataset was produced by combining each of the independently processed data subsets.

### ULM reconstructions of young and aged mouse brains reveal distinct vascular phenotypes

A representative example comparison of a young mouse brain and an aged mouse brain (**Figure 4A**) demonstrate several key observations of the cerebral vasculature. In both cases, the cerebral cortical microvasculature comprises a distinct layer of columnar vessels perpendicular to the cortex, ranging from the brain surface to the subcortical border. Generally, the young cortex demonstrates more densely packed microvasculature with a more orderly structure, whereas the aged brain cortex demonstrates some regional hypo-perfusion and a more disorganized appearance. The hippocampal and thalamic regions generally demonstrate similar levels of overall blood perfusion in both the young and aged brain; however, the older cohort exhibits a more stratified distribution of apparent microvascular velocities with a higher proportion of fast microbubble signal appearing in the large vessels. By contrast, the young mouse hippocampal and thalamic microvessels have a more gradual shift from the fast microbubble velocities in large vasculature to the slower microbubble events in the small vasculature. This is also apparent in a significantly increased right skewness (1.16 vs. 0.98, p = 0.002) of the distribution of blood velocities across the whole depth of the brain in the aged cohort versus the young cohort.

**Figure 4.**
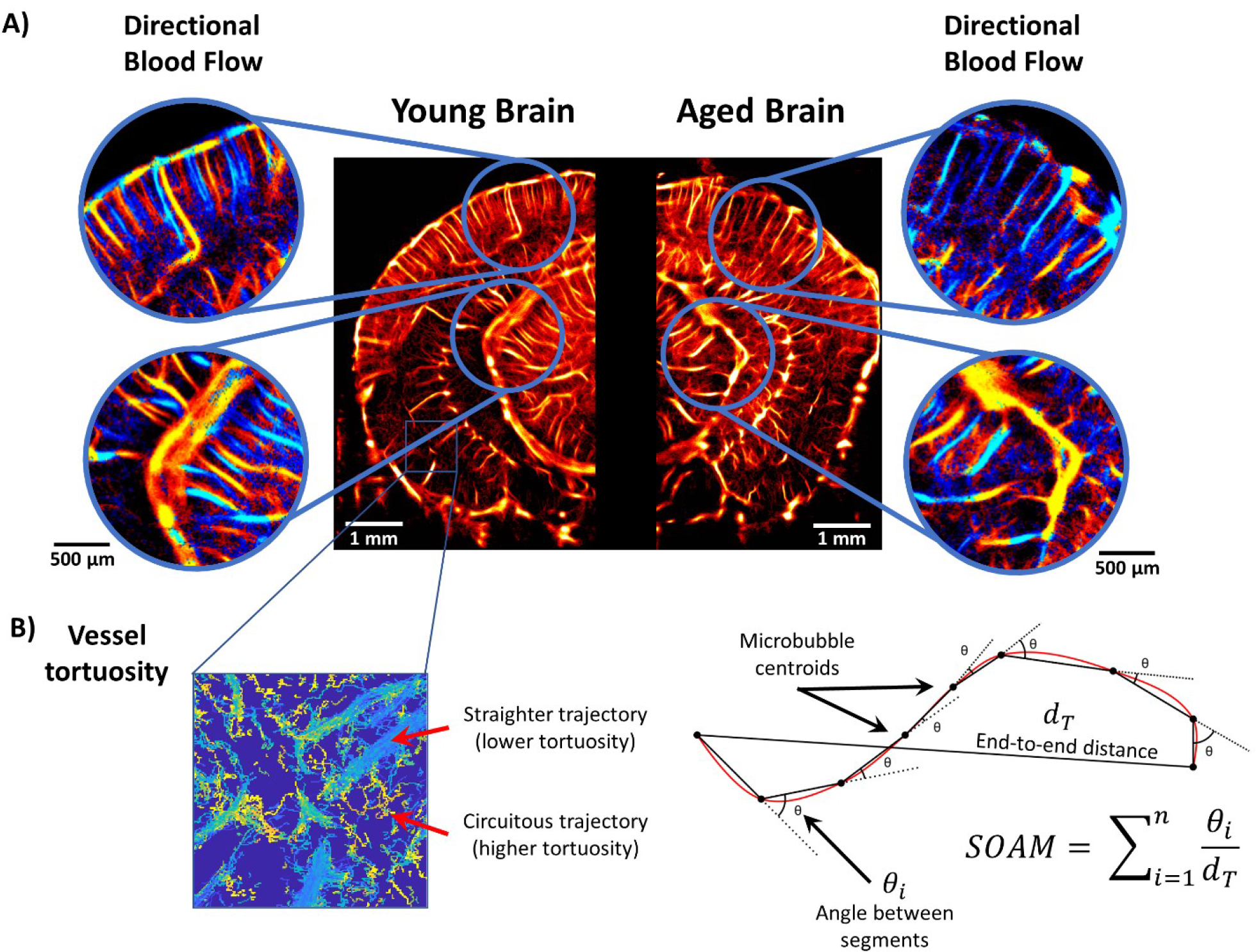
ULM imaging of young and aged mouse brain. (**A**) Example ULM microvessel density maps images from a (left) young mouse brain and (right) an aged mouse brain. Local cortical and subcortical regions are selected and magnified to show the corresponding directional blood flow maps (blue toward the transducer, red away from the transducer). (**B**) An inset image demonstrating distinct vessel tortuosity phenotypes. The larger, straighter vessels have a relatively low measured tortuosity, whereas the smaller connecting vessels are more circuitous. A diagrammatic example of how the sum of angles metric was determined using the microbubble trajectory data. For each complete microbubble trajectory the total path length, end-to-end distance, and angle between every segment was used in the calculation.

### Aged mice have quantitatively different vasculature than young mice

The aged and young mouse cohorts were quantitatively compared based on super-resolution ULM measurements of their auditory cortex, auditory thalamus, entorhinal cortex, hippocampus, superior colliculus, and visual cortex (**Figure 5**). Blood volume was estimated using the mean number of microbubbles that entered a particular ROI. The auditory cortical blood volume (**Figure 5A**) of the aged mouse group was significantly lower than the young mouse group (p < 0.001) and no significant differences were seen in the auditory thalamus or hippocampus. The blood volume was also found to be significantly decreased in the entorhinal cortex (p = 0.007) and the visual cortex (p = 0.001). Similar observations can made about the vascularity of the young brain versus the aged brain (**Figure 5B**), where a significant decrease in vascularity was only found for the auditory cortical (p = 0.018) and visual cortical (p = 0.02) ROIs. Regional blood velocity demonstrated a significant decrease across all of the measured brain regions in the aged mouse group in comparison to the young mouse group (**Figure 5C**). The effect was the most pronounced in the superior colliculus (p < 0.001) and visual cortex (p < 0.001), with proportionally lowered reduction in velocity for the hippocampal, thalamic, auditory cortical, and entorhinal cortical ROIs (p = 0.015, p = 0.003, p = 0.004, and p = 0.001, respectively). The tortuosity of the brain vasculature was using the established metric known as the Sum of Angles Metric (SOAM). A diagrammatic example detailing the metric is demonstrated in **Figure 4B**. All of the measured brain regions had significantly higher metrics of SOAM tortuosity in the aged mouse group in comparison to the young group (**Figure 5D**). Specifically, we found significant SOAM increases in the auditory cortex (p = 0.004), auditory thalamus (p = 0.006), entorhinal cortex (p = 0.001), hippocampus (p = 0.041), superior colliculus (p = 0.002), and visual cortex (p < 0.001). This difference in vessel structural organization mirrors the age-associated decreases in blood velocity that were seen in these anatomical regions.

**Figure 5.**
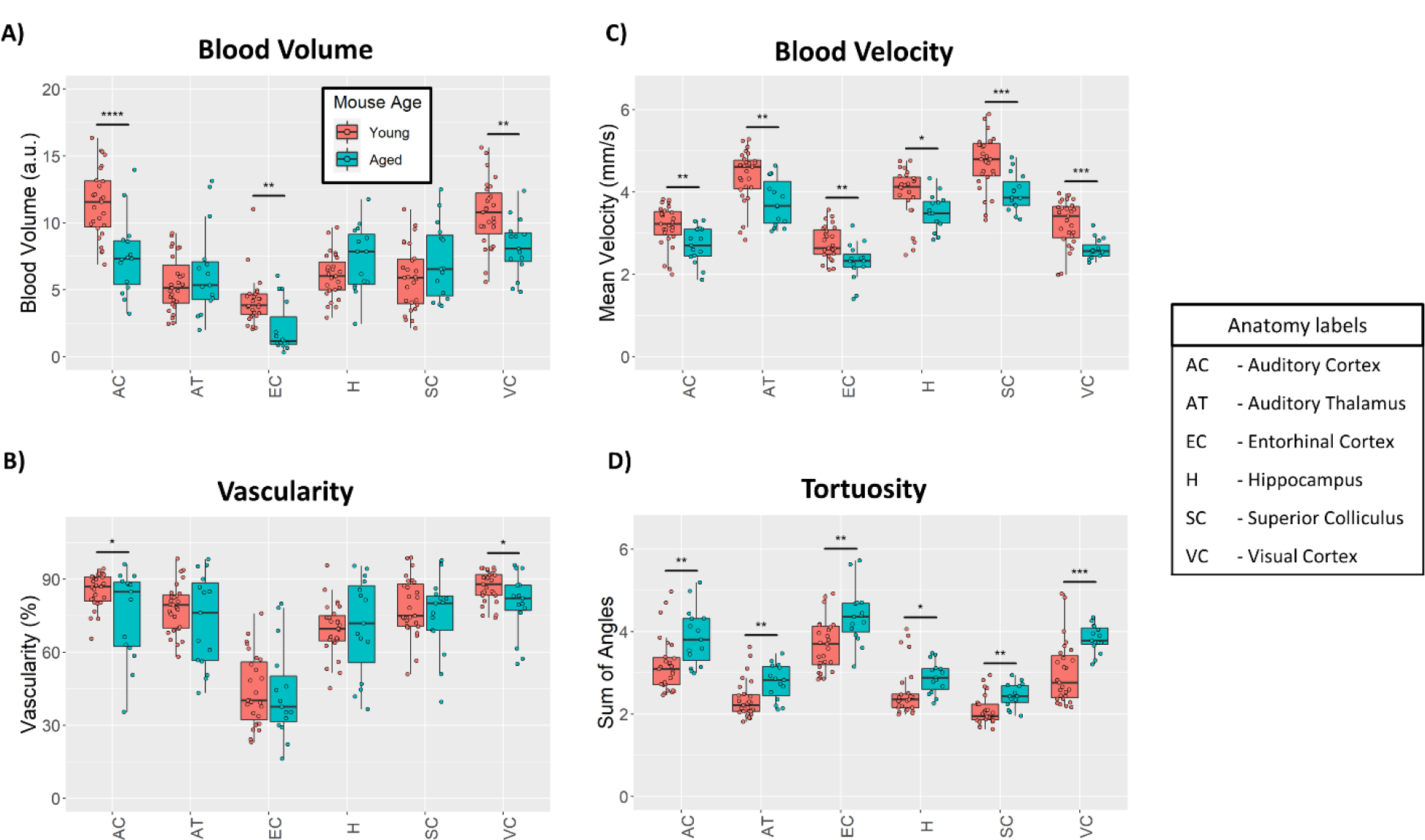
Quantitative ULM measurements. (**A**) Cortical blood volume was significantly decreased in the aged mouse group in comparison to the young group (p < 0.001 for the auditory cortex, p = 0.007 for entorhinal cortex, and p = 0.001 for visual cortex). (**B**) Likewise, ULM measured vascularity demonstrated a significant decrease in the auditory cortex and visual cortex of the brain between the aged and young group (p = 0.018 and p = 0.02, respectively). (**C**) Blood velocity exhibited global decreases across all measured brain regions in the aged mouse cohort. The superior colliculus (p < 0.001) and visual cortex (p < 0.001) demonstrated the most substantial decrease in mean velocity, with proportionally less decrease in the hippocampus (p = 0.015), auditory thalamus (p = 0.003), auditory cortex (p = 0.004), and entorhinal cortex (p = 0.001). (**D**) Vascular tortuosity, as measured by sum of angles metric, showed significant increases across all brain regions (p = 0.004 for auditory cortex, p = 0.006 for auditory thalamus, p = 0.001 for entorhinal cortex, p = 0.041 for hippocampus, p = 0.002 for superior colliculus, and p < 0.001 for the visual cortex)

## Discussion

This study quantified aging-associated changes in the cerebral microvasculature using super-resolution ULM imaging in a mouse model of aging. We demonstrated that ULM imaging provides access to microvessel structural and functional information throughout the entire depth of a coronal section of the brain, allowing for the simultaneous observation of several distinct superficial and deep anatomical brain regions. This ability to conduct global brain imaging is critical for detecting and quantifying the selective vascular vulnerability that is hypothesized to contribute to the heterogeneous decreases in aging-associated cerebral microvascular density (12,22,23).

We found that the aged brain demonstrated decreases in the cortical vascular density and blood volume as measured by ULM imaging (**Figure 4** and **Figure 5**). This finding is consistent with the literature, which reports that aging is associated with decreases in histological measurements of microvascular density throughout the cerebral blood supply (6–12). A more profound effect was observed when comparing the mean blood flow velocity and vascular tortuosity of the aged cohort versus the young mouse group. We also found global decreases in flow velocity and global increases in tortuosity across all analyzed brain regions of the aged mice. This functional decrease in aging-associated cerebral blood flow is a more direct indicator of impaired vascular supply than inferring such impairments from histological findings which indicate reduced microvascular density. Intuitively, the circuitous blood vessel structures (**Figure 4B**) that are measured using the SOAM metric should lead to increased resistance to blood flow and corresponding regional inefficiencies in blood delivery. This measured difference in vessel structural organization is also reflected in the age-associated decreases in blood velocity observed in anatomical regions throughout the brain.

Furthermore, an interesting observation from a direct comparison of super-resolution images of the young and aged mouse brain (**Figure 3**) was the qualitative appearance of a more stratified distribution of microvascular velocities with a higher proportion of the relatively faster microbubble signal appearing in large vessels in the aged mouse brain. This manifested quantitatively as a significantly increased right skewness (1.16 vs. 0.98, p = 0.002) of the distribution of blood velocities of the whole brain in the aged cohort versus the young cohort, with the heavier tail implying a less gradual shift from slow flowing microvessels to faster arterioles. This observation mirrors the findings of Bell and Ball (12) who conducted a morphometric comparison of hippocampal microvessels in aging humans. They found that aging was associated with increases in the diameters of both capillaries and arterioles and, while the density of capillaries decreased, the density of arterioles increased significantly in the aged brain. It can be speculated that this shift in the proportion of vessel diameters in the aging brain should lead to a corresponding shift in blood flow velocities. A more direct comparison examining the distribution of ULM-determined vessel diameters with respect to age would be interesting; however, a substantial limitation of ULM is that the image reconstruction is stochastic. Vessel lumen are gradually filled in with sparse localizations of microbubble trajectories and without *a priori* knowledge of the vessel structure it is difficult to determine the fully perfused vessel diameters. Conventionally, the measurement of vessel sizes in ULM is done through a process of manual selection and segmentation, which is both laborious and prone to bias, and limits the generalizability of the analysis.

This study has some further limitations that should be addressed. ULM imaging is a relatively new technology that is still being optimized and many of the functional output metrics (such as velocity) lack a well-established gold-standard reference to confirm that the reconstructions are informative of the true physiology. We have previously found that ULM imaging in a tumor model correlated with histological measurements of microvascular density and that ULM measurements of tortuosity were informative of tissue hypoxia (33). The choice of anesthesia could also impact the underlying physiology and therefore the quantifications of cerebral blood flow. In this study, we anesthetized the mice for imaging using vaporized isoflurane mixed with oxygen. However, it is well-established that isoflurane has a dose-dependent dilatory effect on the cerebral blood flow (51), which is particularly problematic, as aged mice can become more sensitive to anesthesia. This uncertainty in the cerebral blood supply is also exacerbated by the requirement for long imaging acquisitions times in ULM imaging to ensure that the majority of the microvasculature has been perfused with microbubbles (49,52,53). The ULM imaging protocol in this study typically requires that the mouse be under anesthesia for at least 2 hours total to perform the craniotomy, secure the animal to the imaging stage, place the tail vein catheter, and to acquire the data.

With these caveats in mind, we have demonstrated that ULM imaging can provide subcortical microvascular dynamics *in vivo* to inform vascular changes associated with aging-related cognitive decline. Given the lack of tissue destruction and safety profile of ULM imaging, this technique lends itself well to longitudinal studies of cerebral blood flow to further examine pathological neurodegeneration and to elucidate vasculature-based mechanisms of neuroprotection. Longitudinal study design could either implement a chronic cranial window using an acoustically transparent plastic, such as polymethylpentene (54), or could conduct transcranial imaging with sophisticated phase-aberration correction (55,56). Based on the current findings, a larger study examining the microvascular structure and function of additional cohorts of mouse ages is warranted.

## Author contributions statement

MRL, NCS, DL, and PS designed and wrote the paper. NCS and DL prepared the mouse model and performed craniotomies. MRL, WZ, ZD, and PS performed the ultrasound imaging. ZD designed the ultrasound transducer holder and programmed the motorized imaging stage. MRL and NCS performed image segmentation and image analysis. MRL, XC, and PS developed and applied the super-resolution ULM algorithm.

## Data Availability

The data that support the findings of this study are available from the corresponding authors on request.

## Competing Interests

The authors declare no competing interests.

## Funding

The study was partially supported by the National Cancer Institute, the National Institute of Biomedical Imaging and Bioengineering, the National Institute on Deafness and Other Communication Disorders, and the National Institute on Aging of the National Institutes of Health under grant numbers R00CA214523, R21EB030072, R21DC019473 and R03AG059103. The content is solely the responsibility of the authors and does not necessarily represent the official views of the National Institutes of Health. NCS is supported by a Beckman Institute Postdoctoral Fellowship.

## Notes

### Competing Interest Statement

The authors have declared no competing interest.

